# PRMT5 orchestrates EGFR and AKT networks to activate NFκB and promote EMT

**DOI:** 10.1101/2024.01.03.574104

**Authors:** Lei Huang, Manasa Ravi, Xiao-Ou Zhang, Odette Verdejo-Torres, Noha A.M. Shendy, Mohammad A.M. Nezhady, Sneha Gopalan, Gang Wang, Adam D. Durbin, Thomas G. Fazzio, Qiong Wu

**Affiliations:** Department of Pediatrics, University of Massachusetts Medical School, Worcester, MA, USA; Program in Bioinformatics and Integrative Biology, University of Massachusetts Medical School, Worcester, MA, USA; School of Life Sciences and Technology, Tongji University, Shanghai, China; Department of Molecular, Cell, and Caner Biology, University of Massachusetts Medical School, Worcester, MA, USA; Division of Molecular Oncology, Department of Oncology, St. Jude Children’s Research Hospital, Memphis, Tennessee, USA

**Keywords:** PRMT5, EGFR, AKT, NFκB, EMT, SNAIL, TWIST1

## Abstract

Neuroblastoma remains a formidable challenge in pediatric oncology, representing 15% of cancer-related mortalities in children. Despite advancements in combinatorial and targeted treatments improving survival rates, nearly 50% of patients with high-risk neuroblastoma will ultimately succumb to their disease. Dysregulation of the epithelial-mesenchymal transition (EMT) is a key mechanism of tumor cell dissemination, resulting in metastasis and poor outcomes in many cancers. Our prior work identified PRMT5 as a key regulator of EMT via methylation of AKT at arginine 15, enhancing the expression of EMT-driving transcription factors and facilitating metastasis. Here, we identify that PRMT5 directly regulates the transcription of the epidermal growth factor receptor (EGFR). PRMT5, through independent modulation of the EGFR and AKT pathways, orchestrates the activation of NFκB, resulting in the upregulation of the pro-EMT transcription factors ZEB1, SNAIL, and TWIST1. Notably, EGFR and AKT form a compensatory feedback loop, reinforcing the expression of these EMT transcription factors. Small molecule inhibition of PRMT5 methyltransferase activity disrupts EGFR/AKT signaling, suppresses EMT transcription factor expression and ablates tumor growth *in vivo*. Our findings underscore the pivotal role of PRMT5 in the control of the EMT program in high-risk neuroblastoma.

## Introduction

Neuroblastoma, a complex pediatric malignancy originating from neural crest cells, represents one of the most challenging cancers in pediatric oncology due to its heterogeneous nature and variable clinical outcomes^1–3^. Despite advancements in treatment, high-risk neuroblastoma continues to have a dismal prognosis, underscoring the critical need for novel therapeutic targets and a deeper understanding of its molecular underpinnings^4^.

Protein arginine methyltransferase 5 (PRMT5), a type II arginine methyltransferases member, is the primary methyltransferase generating symmetric dimethylarginine (SDMA)^5, 6^. PRMT5 has emerged as a pivotal regulator of cancer biology and has been implicated in various cancers, including neuroblastoma^7–12^. Our previous research has elucidated a novel role for PRMT5 in modulating AKT phosphorylation through direct methylation, revealing a critical pathway by which PRMT5 impacts tumor growth and metastasis^8^. Furthermore, we demonstrated that PRMT5 regulates expression of the epithelial-mesenchymal transition (EMT) transcription factors such as SNAIL, TWIST1, and ZEB1 via AKT, establishing a link between PRMT5 activity and the EMT process in neuroblastoma^8^. Expression of these transcription factors has recently been linked to a high-risk “mesenchymal” cell state in neuroblastoma, that is relatively resistant to conventional chemotherapies^13–15^, making the processes by which this state is controlled of central importance to the disease^16^.

The AKT pathway, a well-established signaling cascade, plays a crucial role in regulating cell survival, growth, and metabolism and has been extensively studied in the context of various cancers, including neuroblastoma^17, 18^. Epidermal Growth Factor Receptor (EGFR) pathway is another critical signaling mechanism known for its involvement in tumorigenesis and progression in numerous cancers^19^. EGFR is overexpressed in neuroblastoma patients and is associated with resistance to ALK inhibitors and chemotherapy^20, 21^. Further, exogenous EGF supplementation of neuroblastoma cells *in vitro* can partially promote a more mesenchymal phenotype^22^. However, how EGFR functions to promote these transitions in neuroblastoma has not been thoroughly investigated, and mechanisms to disrupt this for clinical benefit have remained as of yet, unexplored. Further, the interrelation and combination of the impact of AKT and the EGFR pathway on EMT, a key process in tumor metastasis and progression, remains less explored in this context.

Here, we reported that PRMT5 regulates EGFR transcription and AKT activation independently. Importantly, EFGR and AKT form a compensatory feedback loop to reinforce EMT, although they cannot compensate for deficiency of the opposite protein in the setting of PRMT5 inhibition. We identify NFkB as the effector downstream of AKT/EGFR that promotes the expression of the pro-EMT transcription factors SNAIL, TWIST1, and ZEB1, and demonstrate that disruption of PRMT5 activity ablates these signaling pathways, EMT transcription factor expression and tumor growth, *in vivo*. These findings have significant implications for understanding the molecular landscape of neuroblastoma and open new avenues for therapeutic interventions targeting the PRMT5-AKT-EGFR-NFkB network in high-risk, mesenchymal subtype neuroblastoma cells.

## Results

### PRMT5 expression levels negatively correlate with NB patient survival

To explore the effects of PRMT5 expression with patient prognosis in neuroblastoma, we performed a correlational analysis of PRMT5 expression with patient prognosis using RNA-seq datasets from three independent and non-overlapping patient cohorts (Kocak, SEQC, and Cangelosi) from the R2 database (R2: Genomics Analysis and Visualization Platform [http://r2.amc.nl]). Patients with the highest quartile of PRMT5 expression exhibited a significantly worse prognosis than those with the bottom quartile in all datasets tested (Fig. 1a-c). Since these datasets include a variety of patients with neuroblastoma, ranging from low to high-risk, we next sought to understand the effect of PRMT5 expression levels in only the highest risk patients. When focused only on stage 3 and 4 neuroblastoma patients from the SEQC dataset, we observed a striking finding: approximately 70% of patients with the highest quartile of PRMT5 transcript levels succumbed to the disease within a 4-year timeframe, in contrast to patients in the lowest quartile, who remained alive for an extended follow-up period of over 192 months (Fig. 1d). These results strongly link the expression of PRMT5 to patient outcome in high-risk neuroblastoma.

**Figure 1.**
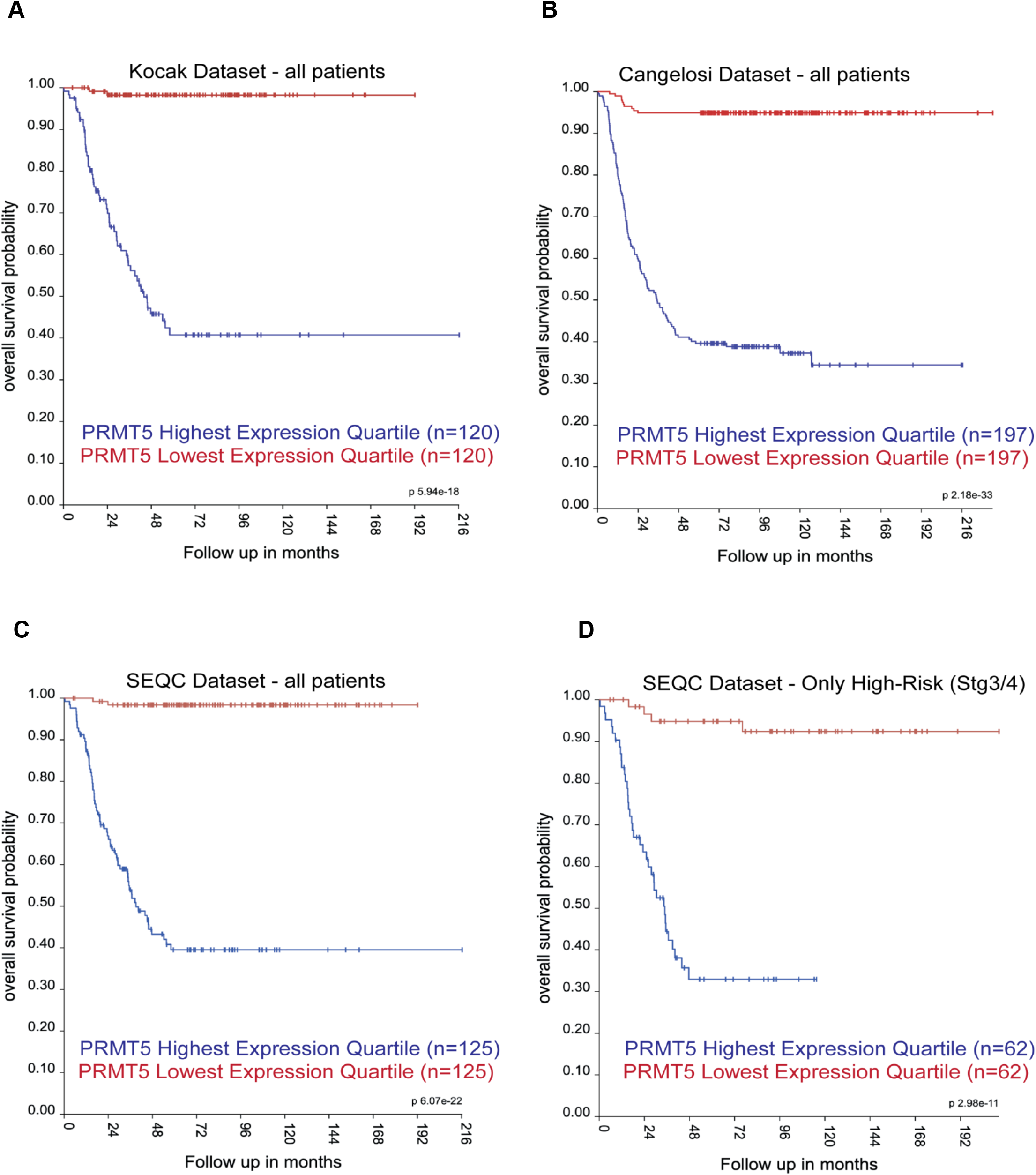
PRMT5 expression levels negatively correlated to neuroblastoma patient survival. Overall survival is mapped using the top and bottom quartile of PRMT5 expression in all patients from the Koca dataset (**a**), Cangelosi dataset (**b**), and SEQC dataset (**c**). Additionally, overall survival of high-risk patients (stage 3/4) in the SEQC dataset was analyzed (**d**). Original data was extracted from www.r2.amc.nl.

### Global transcriptome analysis revealed multiple PRMT5-dependent pathways in neuroblastoma

Next, we sought to understand the mechanisms by which PRMT5 regulates neuroblastoma gene expression, using RNA-seq. To do so, we used examples of *MYCN-*amplified (NGP) and non-amplified (CHLA20) cells, which represent distinct subgroups of high-risk clinical disease. Changes in global gene expression were identified in CHLA20 and NGP cells treated with DMSO or the potent and specific PRMT5 methyltransferase inhibitor, GSK591 by unsupervised hierarchical clustering (Fig. 2a). There was a small subset of overlap in the differentially expressed genes (DEGs) in these two cell lines, which may indicate the different gene networks mediated by expression of the driving oncogenes *c-MYC* (in CHLA20) compared with *MYCN* (in NGP), respectively (Fig. 2b). By performing a KEGG (Kyoto Encyclopedia of Gene and Genomes) pathway analysis, we observed that downregulated DEGs in both cell lines were commonly enriched in PI3K-AKT and p53 signaling pathways(Fig. 2c-d), supporting our previous finding^8^. These results were confirmed by performing gene set enrichment analysis, which demonstrated that GSK591-treated CHLA20 cells showed reduced AKT up-and down-regulated genes (Fig. 2e), and similarly, in NGP cells, GSK591 treatment downregulated AKT/mTOR specific genes (Fig. 2f).

**Figure 2.**
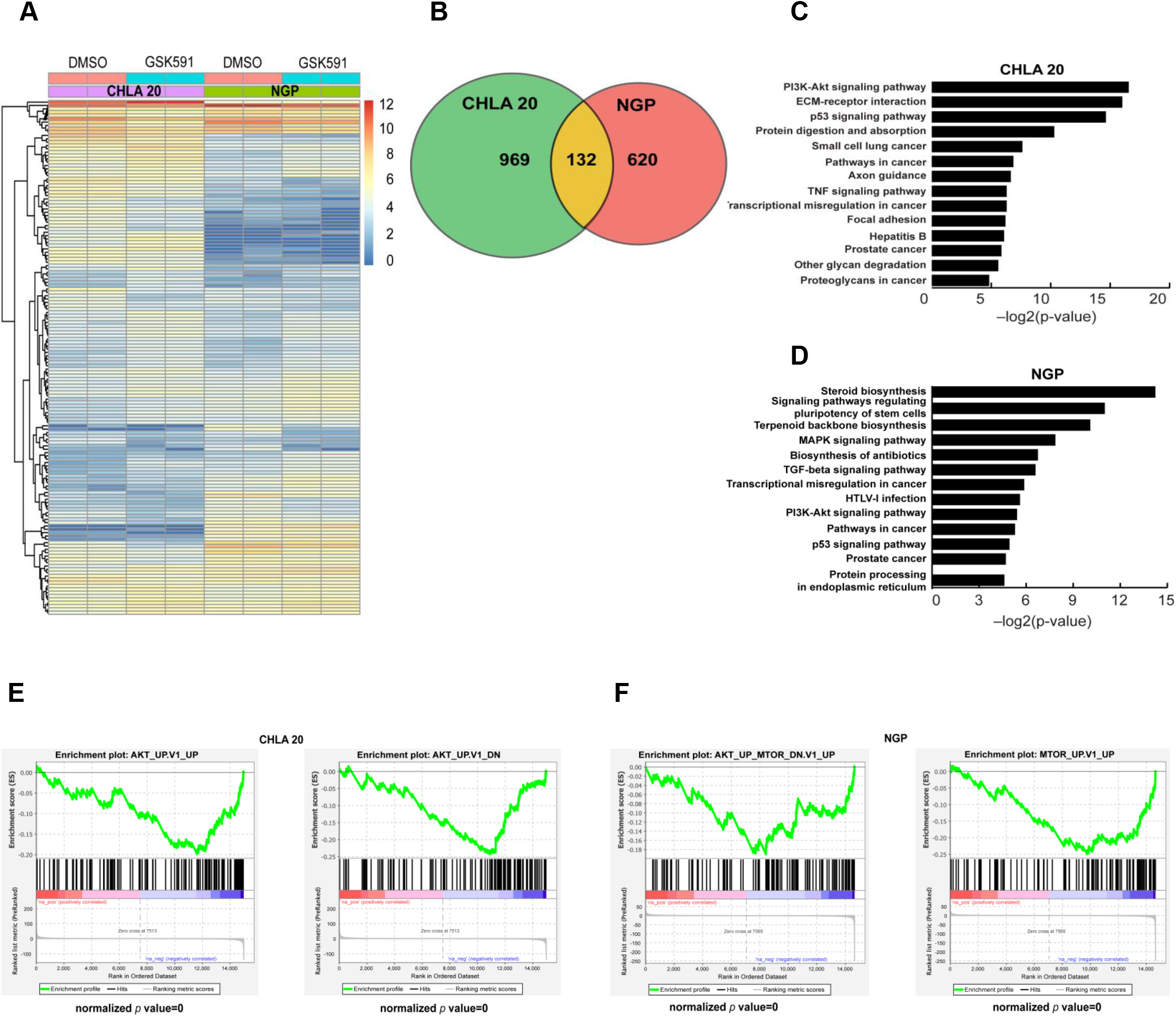
Global transcriptome analysis of PRMT5 inhibition in neuroblastoma cells. **a**, Heat map of significantly differentially expressed genes (fold change >2 and FDR<0.01) between cells treated with DMSO or GSK591. The color scale represented log2 normalized gene expression in DMSO and GSK591 treatment (n = 2 biologically independent samples). **b**, Venn diagram of genes differentially expressed under GSK591 treatment in CHLA20 and NGP (p=7.736×10-48, hypergeometric test). Enriched Kyoto Encyclopedia of Genes and Genomes (KEGG) pathways analysis in CHLA 20 (**c**) and NGP under GSK591 treatment (**d**). Gene set enrichment analysis of AKT signaling signature genes in CHLA20 cells treated with GSK591 (**e**) and AKT-mTOR signaling signature genes in NGP cells treated with GSK591 (**f**).

### PRMT5 inhibition impairs EGFR signaling

Receptor tyrosine kinases (RTKs) regulate pivotal signaling pathways in cancer, such as SRC proto-oncogene, epidermal growth factor receptor (EGFR), fibroblast growth factor receptor (FGFR), and human epidermal growth factor receptor 2 (HER2)^23, 24^. Analysis of our RNAseq data demonstrated that specific receptors, including *EGFR*, *fibroblast growth factor receptor 4* (*FGFR4*), and *neural growth factor receptor (NGFR)* were all downregulated by GSK591 treatment, which we confirmed by RT-qPCR (Fig. 3a). Further, we also observed that either chemical inhibition of PRMT5 using GSK591 (Fig. 3b), or shRNA-depletion of PRMT5 (Fig. 3c) also resulted in downregulation of EGFR, FGFR4 and NGFR by western blotting. Of note, EGFR is overexpressed in neuroblastoma patients and associated with resistance to ALK inhibitors and chemotherapy^20, 21, 25^, and supplementation of EGF to neuroblastoma cell lines causes changes in gene expression enforcing a partial mesenchymal state^22^, making this a target of exceptional interest in neuroblastoma. Therefore, we examined EGFR signaling in cells treated with increasing doses of GSK591. As shown in Fig. 3d, the compound drastically decreased global cellular phospho-tyrosine modifications in both NGP and CHLA20 cells, in a dose-dependent fashion. Moreover, GSK591-treatment severely blunted the neuroblastoma cellular response to exogenous EGF stimulation, indicating that PRMT5 inhibition impairs EGFR signaling (Fig. 3e). We previously reported that a group of well-known cancer associated RTKs, including IGF1R, VEGFR, and ERBB3, were not affected by GSK591 treatment at either the protein expression or phosphorylation levels^8^. In contrast, these results also suggested that PRMT5 may control EGFR signaling in neuroblastoma. In addition to its effects on AKT, PRMT5 also performs methylation of arginine residues on Histones H3 and H4, thereby affecting gene transcription^26–28^. Thus, we next explored whether PRMT5 proteins were localized at the *EGFR* promoter. Chromatin immunoprecipitation (ChIP)-PCR experiments identified enrichment of PRMT5 at the *EGFR* promoter, concurrent with the presence of the PRMT5-catalyzed histone marks, H3R8 and H4R3 symmetric di-methylarginine (SDMA) and RNA polymerase II (Fig. 3f). Importantly, these associations were diminished in PRMT5 knockdown cells, suggesting that PRMT5 binding is required for RNA polymerase II presence at the *EGFR* promoter (Fig. 3f). Further, while catalytic inhibition of PRMT5 with GSK591 did not affect the association of PRMT5 with the *EGFR* promoter, it did reduce SDMA on H3R8 and H4R3, as well as Pol II binding at this locus (Fig. 3f). These results demonstrate that PRMT5 enzymatic activity is responsible for the epigenetic activation of EGFR expression.

**Figure 3.**
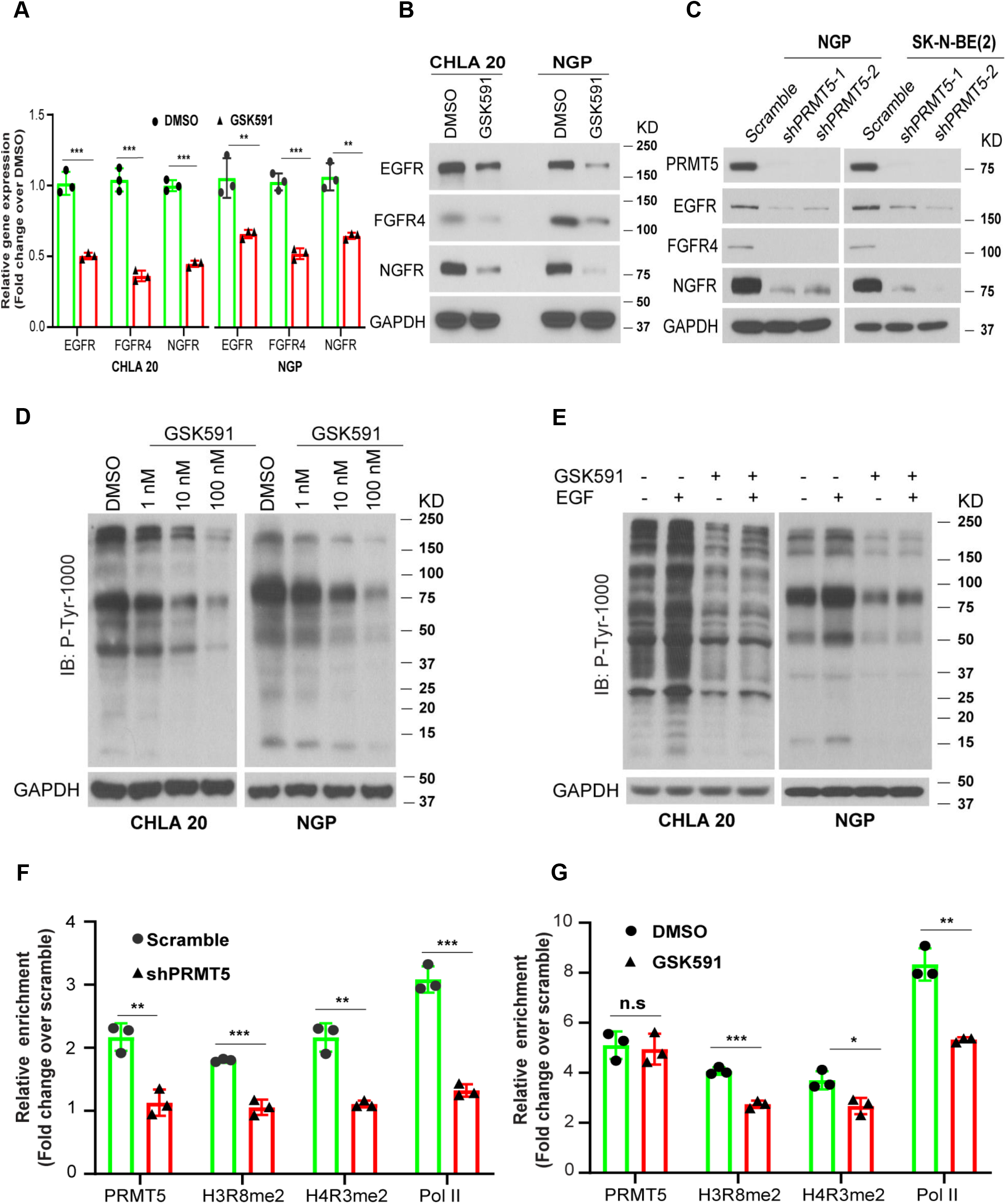
PRMT5 inhibition impairs EGFR signaling. Neuroblastoma cells were treated with DMSO or 100 nM GSK591 for six days. **a**, RT-qPCR analysis of EGFR, FGFR4, and NGFR expression in CHLA20 and NGP cells treated with DMSO or GSK591. **b**, EGFR, FGFR4, and NGFR protein levels in neuroblastoma cells treated with DMSO or GSK591. **c**, Western blots of EGFR, FGFR4, and NGFR in control and PRMT5 knockdown cells. **d**, EGFR signaling examined by immunoblotting with Phospho-Tyrosine (P-Tyr-1000) MultiMab™ Rabbit mAb mix in cells treated with DMSO or increasing doses of GSK591. GAPDH was used as a loading control. **e**, The response to EGF stimulation measured by P-Tyr-1000 on cells treated with DMSO or GSK591. The occupancy of PRMT5, H3R8mes2, H4R3mes2, and Pol II at EGFR promoter measured by chromatin immunoprecipitation (ChIP) in NGP cells harboring scramble or shPRMT5 (**f**), and CHLA20 cells treated with DMSO or GSK591 (**g**). **a**-**g**, n=3. Data were analyzed by multiple *t*-test in GraphPad Prism, \**p* < 0.05; \*\**p* < 0.01; *** *p* < 0.001.

### EGFR and AKT are independent downstream targets of PRMT5

Previously, we demonstrated that PRMT5 controls activation of AKT1 in neuroblastoma^8^. Although PRMT5 inhibition reduced gene expression and protein levels of EGFR, overexpression of EGFR was not sufficient to rescue the decreased AKT phosphorylation^8^. These observations suggested that PRMT5 may control AKT and EGFR independently. To further delineate the structure and interactions of this regulatory circuit of PRMT5, EGFR, and AKT, we re-expressed EGFR, FGFR4, or NGFR in neuroblastoma cells treated with GSK591. Here, we used CHLA20 *MYCN-*non-amplified and a separate *MYCN-*amplified neuroblastoma cell line, SK-N-BE2(c). As shown in Fig. 4a, exogenous re-expression of these RTKs was not sufficient to rescue the decreased phospho-AKT levels in GSK591-treated cells. AKT has been shown to facilitate the trafficking and recycling of EGFR^29^. To determine whether the impaired AKT activation also contributes to the reduction of EGFR protein levels, we treated neuroblastoma cells with direct AKT kinase inhibitor MK-2206 alone or in combination with GSK591. MK-2206 treatment blocked AKT phosphorylation in control and GSK591 treated cells, indicating the effectiveness of MK-2206 at the doses used (Fig. 4b,c). However, the addition of MK-2206 failed to further affect protein levels of EGFR or PRMT5, nor the phosphorylation of EGFR, driven by GSK591 treatment (Fig. 4b,c). Moreover, treatment of GSK591, even at very low doses (1-10 nM), was sufficient to enhance the effects of the EGFR inhibitor Erlotinib on cell viability in CHLA20 and NGP cells (Fig. 4d). Thus, these results demonstrate that PRMT5 regulates AKT and EGFR in a direct and independent manner.

**Figure 4.**
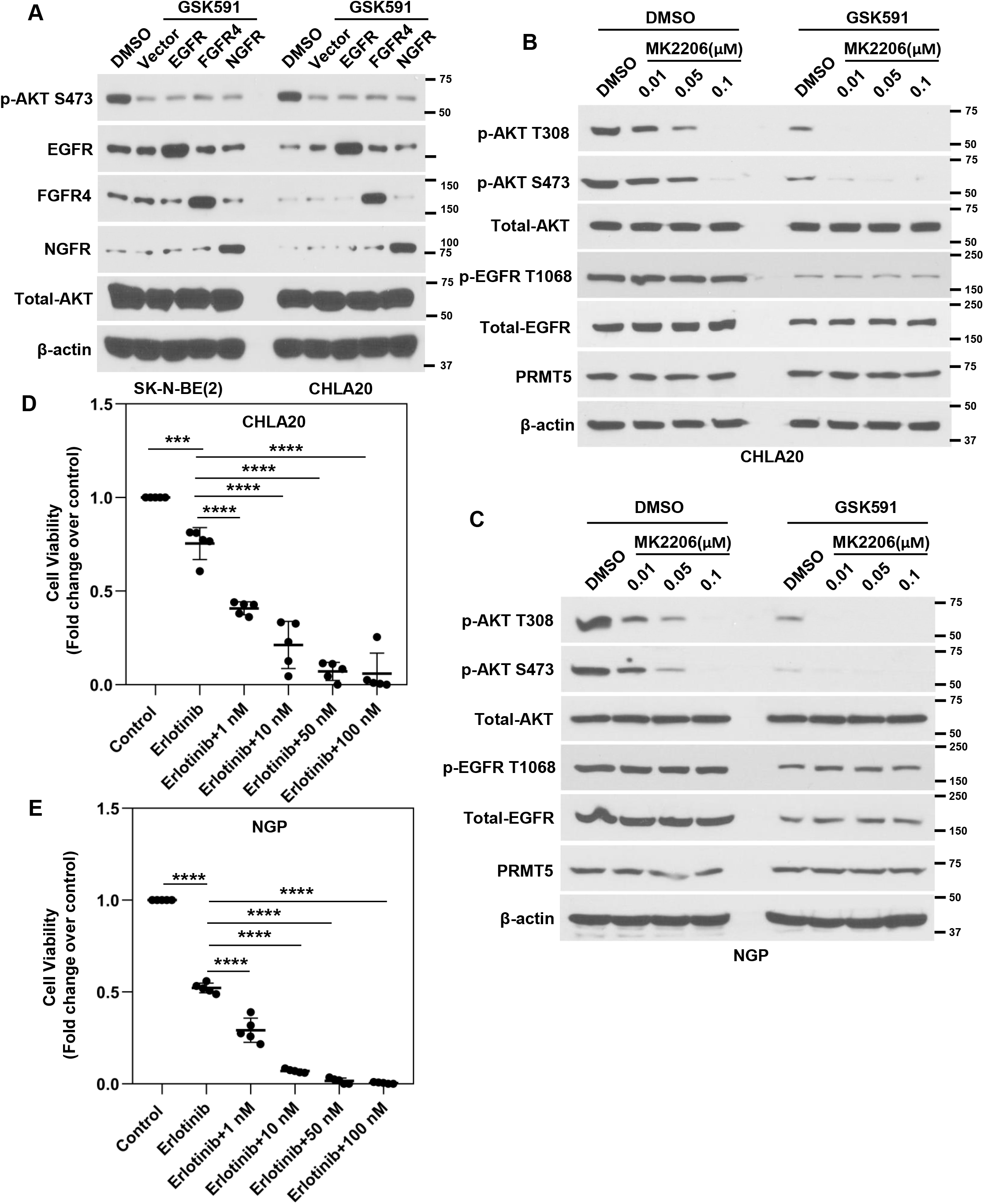
EGFR and AKT are independent downstream targets of PRMT5. **a,** phosphorylation of AKT was examined in GSK591-treated NB cells with exogenous EGFR, FGFR4, and NGFR. **b**, the total protein levels and phosphorylation of EGFR in NB cells treated with DMSO or GSK591 in the presence of MK-2206. **c**, cell viability was examined by MTS assay in NB cells treated with Erlotinib with increasing doses of GSK591. Data were analyzed by multiple *t*-test in GraphPad Prism, *** *p* < 0.001, **** *p* < 0.0001.

### EGFR and AKT form a complementary loop to support EMT

Previously, we demonstrated that PRMT5-mediated AKT1 activation controls EMT program in neuroblastoma^8^. EGFR signaling is a critical transduction mechanisms underlying the induction of SNAIL-dependent EMT in other tumor models^30–32^, and is associated with a more mesenchymal state in neuroblastoma^33^. Thus, we sought to determine whether the reduced EGFR expression and activity, driven by PRMT5, also contributed to this phenomenon independent of AKT activity. Indeed, the protein levels of pro-EMT-transcription factors (EMT-TFs) such as ZEB, TWIST1, and SNAIL were downregulated in cells treated with the EGFR inhibitor, erlotinib (Fig. 5a). Moreover, exogenous expression of EGFR increased the protein levels of these EMT-TFs (Fig. 5b). As a phenotypic outcome of mesenchymal cell state, we observed that cell invasion through the extracellular matrix was severely attenuated under erlotinib treatment, whereas overexpressing EGFR enhanced cell invasion in CHLA20 (Fig. 5c-d) and NGP (Fig. 5e-f). These results suggested that EGFR signaling plays a role in regulating the EMT program in neuroblastoma cells.

**Figure 5.**
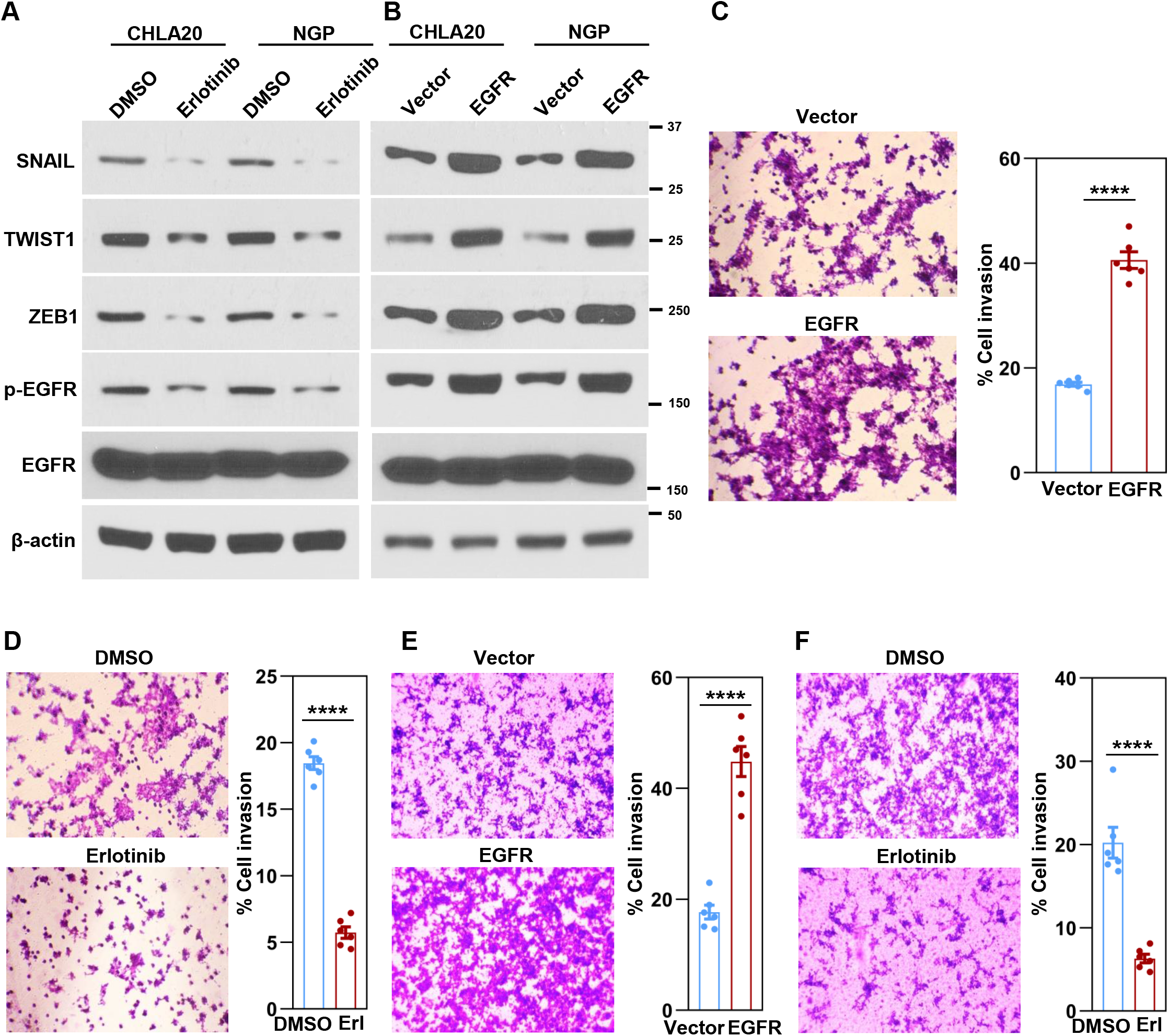
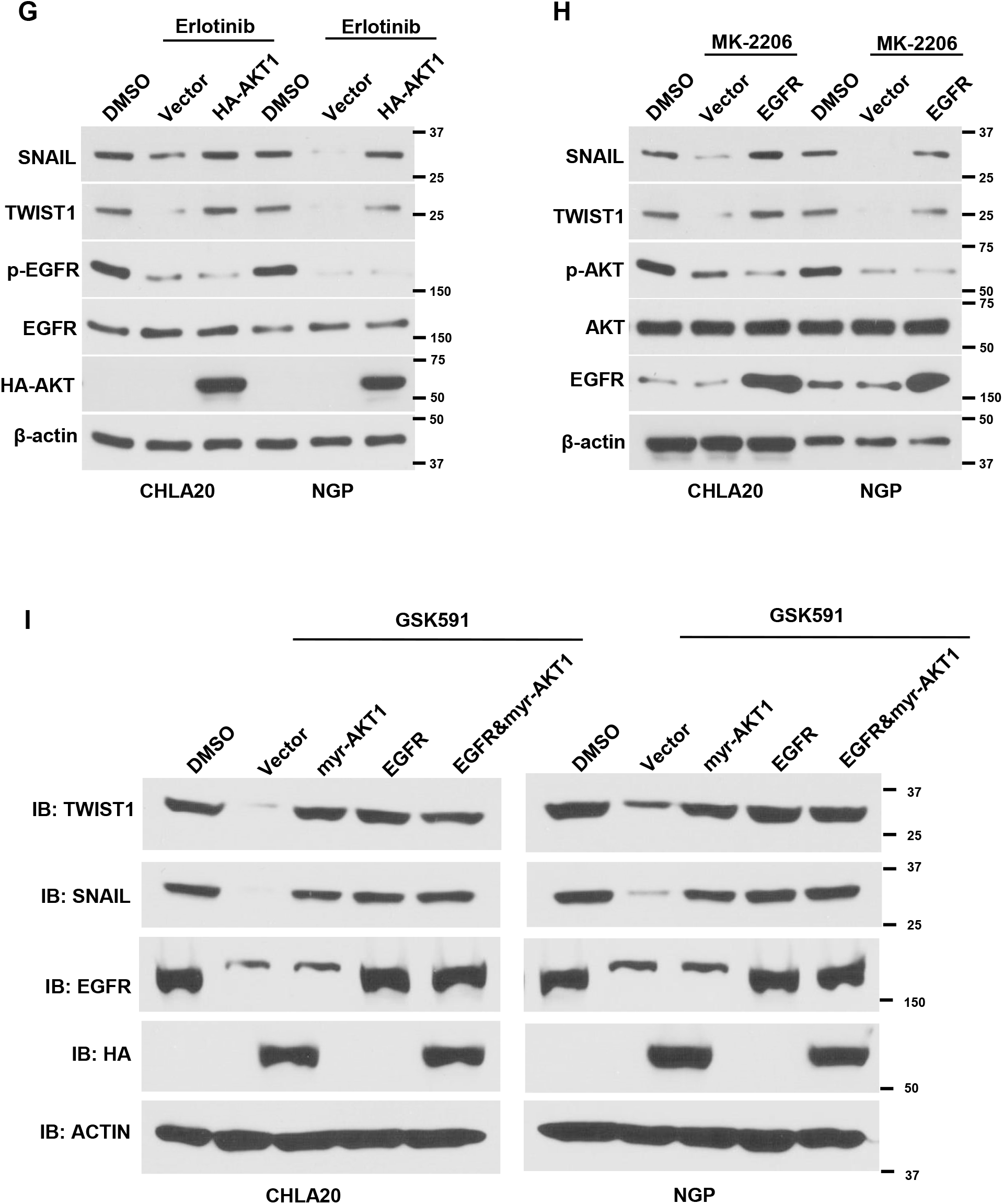
EGFR and AKT form a complementary loop to support EMT. SNAIL, TWIST1, and ZEB1 protein levels in NB cells treated with Erlotinib (**a**) or transfected with a control vector or EGFR (**b**). Cell invasion capacity was examined by Matrigel transwell invasion assay in CHLA20 either in the presence of Erlotinib (**c**) or exogenous EGFR (**d**) as well as in NGP cells either in the presence of Erlotinib (**e**) or exogenous EGFR (**f**). Representative images were shown on the left, scale bars, 100 μm. The percentage of invasive cells normalized by cell numbers in the non-ECM-coated 12-well plate using Image J (n=6) was shown on the right. Data were analyzed by two-tail *t*-test in GraphPad Prism, **** *p* < 0.0001. **g**, SNAIL, TWIST1, and ZEB1 expression in DMSO or Erlotinib-treated NB cells with or without exogenous AKT1. **h**, SNAIL and TWIST1 expression levels were examined in DMSO or MK-2206 treated NB cells with or without exogenous EGFR. **i**, the protein levels of SNAIL and TWIST1 in DMSO or GSK591-treated NB cells transfected vector, myr-AKT1, EGFR, or both.

Next, we sought to understand the interactions between AKT and EGFR, in regulation of EMT-TF expression. First, we established cells overexpressing HA-AKT1, and examined the effects of erlotinib treatment. In these cells, we observed that overexpressed HA-AKT1 was sufficient to rescue the decreased protein levels of TWIST and SNAIL (Fig. 5g). Similarly, exogenous EGFR expression was able to restore the expression of SNAIL and TWIST in MK-2206 treated cells (Fig. 5h). These data suggested that EGFR and AKT are able to function in parallel pathways to control expression of the EMT-TFs TWIST1 and SNAIL. Since PRMT5 inhibition with GSK591 causes loss of EGFR expression and AKT phosphorylation, we next examined the effect of overexpressing EGFR or constitutively activated myr-AKT1 in GSK591 treated cells. In this setting, overexpression of either EGFR or myr-AKT1 alone was sufficient to rescue the protein levels of TWIST1 and SNAIL (Fig. 5i). Further, combination of expression of EGFR or myr-AKT1 had no increased effect on expression of TWIST1 and SNAIL (Fig. 5i) suggestive that each individual protein was sufficient to drive maximal expression. Intriguingly, although FGFR4 and NGFR levels were also reduced by GSK591, forced expression of these proteins had no impact on reduced EMT transcription factors (Supplemental Fig. 1). These results suggested that EGFR and AKT form a compensatory feedback loop reinforcing the expression of these EMT transcription factors and suggest that targeting EGFR or AKT alone may be inadequate to block EMT and metastasis.

### PRMT5-dependent EGFR and AKT signalings regulate NFκB activation to promote EMT

NFκB has been reported to regulate EMT in various types of cancers such as breast cancer, lung cancer, and pancreatic cancer^34–36^. Moreover, NFκB was shown as a downstream effector of multiple signaling pathways, including EGFR and AKT^37–39^. Thus, we hypothesized that EGFR and AKT may converge on NFκB to regulate the EMT program in an independent and complementary manner. Indeed, treatment of GSK591, MK-2206, or Erlotinib all reduced the phosphorylation of the p65 subunit of NFκB (Fig. 6a). To further delineate these signaling cascades in regulating EMT, we determined whether PRMT5 expression could specifically regulate NFκB phosphorylation. Overexpressing wild type PRMT5, but not an enzymatically deficient mutant form of PRMT5, increased the phosphorylation of p65-NFκB (Fig. 6b). Similar results on phosphorylated-p65 NFκB were observed with enforced expression of exogenous EGFR or AKT in CHLA20 and NGP cells (Fig. 6b). Notably, in GSK591 treated cells, restoring the decreased AKT activation by a constitutively activated form of AKT (myr-AKT), or reduced EGFR levels by exogenous EGFR, only partially rescued p65-NFκB phosphorylation, suggesting that combined EGFR and AKT activities were required to stimulate maximal p65-NFκB phosphorylation (Fig. 6c). To verify the involvement of NFκB in regulating EMT, we treated NB cells with Bay 11-7082, a small molecule inhibitor that blocks IKKB phosphorylation of p65-NFκB^40^. As shown in Fig. 6d, treatment of Bay 11-7082 markedly reduced p65-NFκB phosphorylation, concomitantly with the decrease of ZEB1, SNAIL, and TWIST levels. These results suggest that NFκB activation is downstream of AKT/EGFR signaling, and critically affected by PRMT5 enzymatic activity.

**Figure 6.**
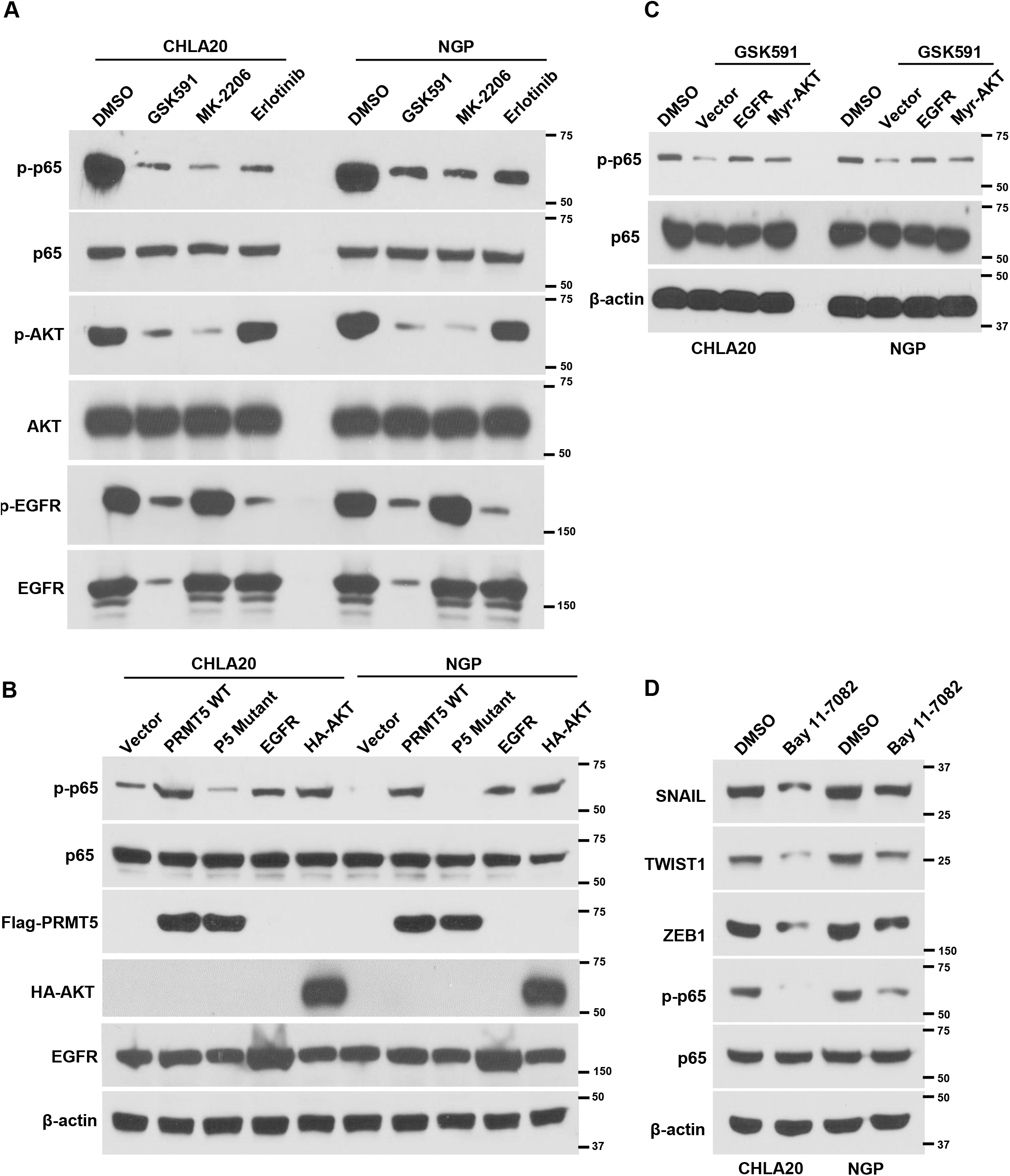
PRMT5-dependent EGFR and AKT signalings regulate NFκB activation to promote EMT. **a**, the total protein level and phosphorylation of NFĸB p65, AKT, and EGFR in NB cells treated with DMSO or GSK591, MK-2206, and Erlotinib. **b**, the expression and phosphorylation of p65 in NB cells transfected with vector, PRMT5 wild type or enzymatically deficient mutant, EGFR, and AKT. **c**, phosphorylation of p65 in DMSO or GSK591-treated cells transfected with vector, EGFR, or a constitutively activated form of AKT. **d**, SNAIL, TWIST1, and ZEB1 protein levels in NB cells treated with NFĸB inhibitor Bay 11-7082.

Since PRMT5 enzymatic activity was required for NFκB activity, and PRMT5 was previously reported to dimethylate R30 or R174 of the p65 subunit to activate NFκB in response to IL-1β and TNF and IFN-γ costimulation, respectively^41, 42^, we next sought to determine if PRMT5 could directly methylate NFκB to affect its phosphorylation. To do so, we performed immunoprecipitations of the p65 subunit of NFκB from DMSO or GSK591 treated NGP or CHLA20 cells, and then examined the SDMA of p65 by Western blot. Although global SDMA was significantly decreased in GSK591 treated cells, the SDMA of p65 was not affected by GSK591 treatment, suggesting other type II PRMT enzymes but not PRMT5, are responsible for methylation of p65 in the context of neuroblastoma (Supplemental Fig. 2). Taken together, these results suggest that PRMT5 regulates NFκB activation via independent regulation of an AKT/EGFR axis and not by direct methylation.

### PRMT5 inhibition reduces tumor growth and prolongs survival in a MYCN-amplified NB PDX model

We previously reported that PRMT5 inhibition or depletion attenuated the growth of SK-N-BE2(c) neuroblastoma cell line xenograft and blocked tumor cell metastasis^8^. To expand upon these results, we sought to understand whether PRMT5 inhibition could be effective on survival and tumor growth in a more physiologically relevant, chemoresistant model. Thus, we next tested the clinical PRMT5 inhibitor GSK595 in a patient-derived xenograft model COG-N-623x. This is a highly chemoresistant PDX derived from a patient with *MYCN*-amplified neuroblastoma who displayed progressive disease despite induction chemotherapy and surgical management, on the Children’s Oncology Group trial ANBL1221. COG-N-623x cells were implanted subcutaneously in NSG mice, and tumor size was measured 3 times a week by digital caliper. Once tumor volume reached stable engraftment and growth at 300-500 mm^3^, mice were randomized to vehicle or GSK595 treatment with 50 mg/kg GSK595 or equal amount of vehicle twice daily by oral gavage as previously^8^. reached close to 2000 mm^3^, and the endpoint of the experiment was set when the last animal in the vehicle group was sacrificed. As shown in Fig. 7a, in contrast to vehicle treated animals which grew rapidly requiring sacrifice within 2-3 weeks, GSK595 treatment dramatically reduced tumor growth, with barely any residual tumor growth detected. Analysis of survival data demonstrated that all tumors in the vehicle group reached 2000 mm^3^ within 3 weeks, however, all animals in the GSK595 group were event-free, with minimally detectable tumors at this timepoint (Fig. 7b). Next, we examined the effects of GSK595 treatment on global SDMA, the phosphorylation of AKT downstream targets, and tyrosine phosphorylated proteins in tumor cells isolated from vehicle or GSK595 treated PDX tumors. To do so, we isolated human tumor cells from PDX tumors and performed western blotting, using antibodies against SDMA, the Phospho- (Ser/Thr) Akt Substrate Motif (RXXS*/T*), or phosphorylated tyrosine residues. In contrast to vehicle treated controls, and consistent with our *in vitro* findings, GSK595-treated tumors displayed strikingly lower levels of these post-translational modifications (Fig. 7c). Further, we performed western blotting to examine the phosphorylation of AKT, p65, and total levels of EGFR in these tumor samples, which displayed a marked reduction in GSK595-treated tumors, with associated decreases in the expression of the EMT-TFs SNAIL, TWIST1, and ZEB1 (Fig. 7d). These results support that PRMT5 inhibition attenuates PDX tumor growth and extends survival in a highly chemoresistant *MYCN-*amplified neuroblastoma PDX, associated with regulation of AKT/EGFR signaling to modulate NFκB activation with suppression of EMT-TFs.

**Figure 7.**
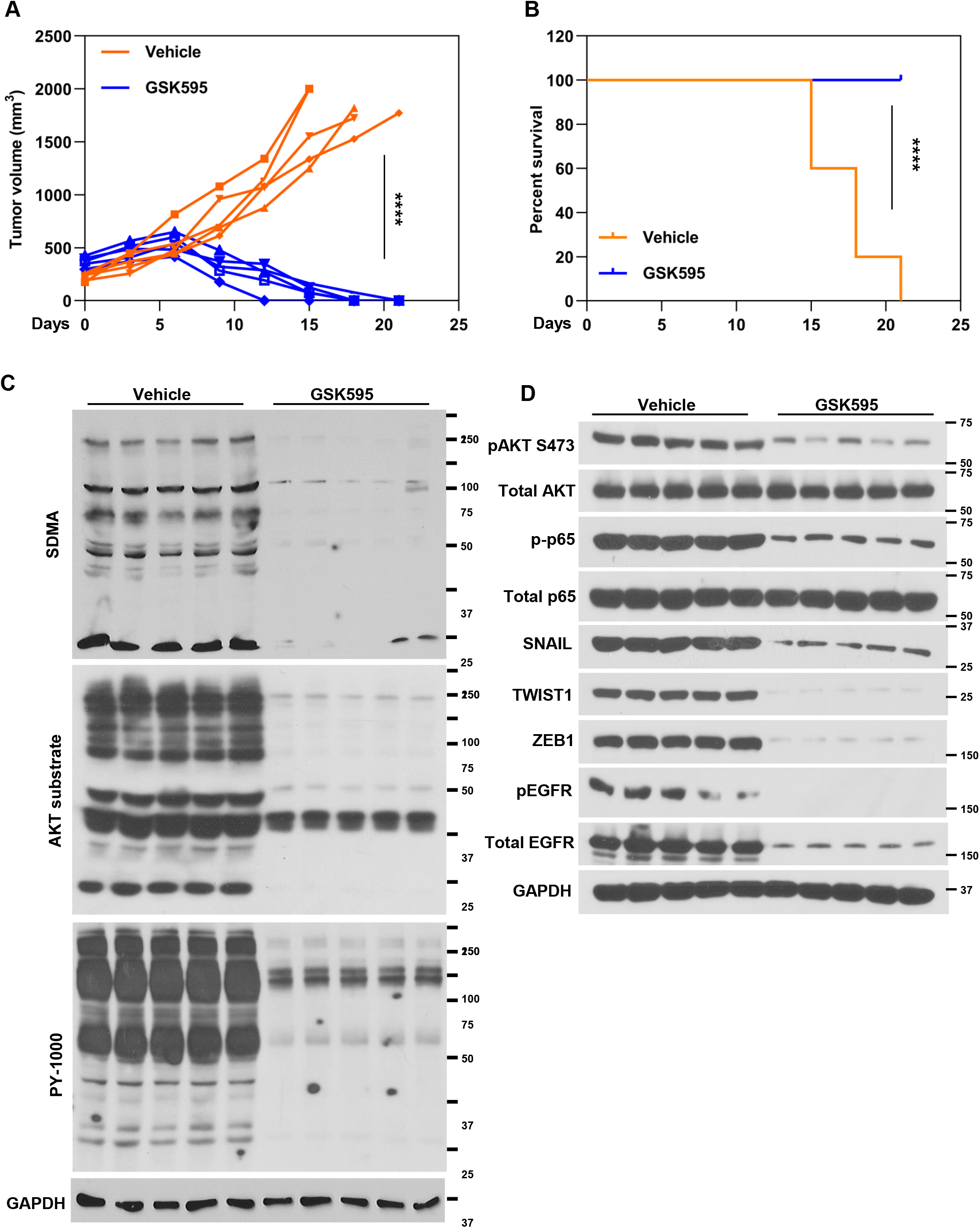
PRMT5 inhibition reduces tumor growth and prolongs survival in a MYCN-amplified NB PDX model. **a** tumor growth curve of the vehicle or GSK595-treated COG-N-623x (n=5). **b**, event-free survival (EFS) of the vehicle or GSK595-treated COG-N-623x (n=5). Data were analyzed by two-tail *t*-test in GraphPad Prism, **** *p* < 0.0001. **c**, phosphorylation of tyrosine proteins and AKT substrates, and the symmetric dimethylarginine proteins in tumor cell lysate from the vehicle or GSK595-treated COG-N-623x. **d**, the phosphorylated AKT, EGFR, and p65 levels, and SNAIL, TWIST1, and ZEB1 expression in lysate extracted from the vehicle or GSK595-treated COG-N-623x tumors.

## Discussion

Here, we showed that PRMT5 is associated with neuroblastoma survival. Further, we demonstrate that PRMT5 regulates the transcription of EGFR in neuroblastoma, which functions as an upstream regulator of the EMT transcription factors SNAIL, TWIST1, and ZEB1. In combination with prior work, these results demonstrate a parallel pathway for PRMT5 activity, in addition to regulation of AKT, to promote EMT in neuroblastoma. Notably, in PRMT5 inhibitor treated neuroblastoma cells, overexpression of EGFR failed to rescue the decreased phosphorylated AKT, nor did myr-AKT restore the reduced protein level of EGFR, indicating that AKT and EGFR are independent downstream targets of PRMT5. It is noteworthy that overexpressing EGFR or constitutively activated myr-AKT alone is sufficient to restore the decrease of EMT-TFs, suggesting that AKT and EGFR form a complementary strategy to reinforce EMT through the control of EMT-TF expression. Furthermore, we showed that NFkB is the downstream effector of AKT and EGFR in regulating EMT. Thus, PRMT5 inhibition represents a strategy for inactivating both of these arms of EMT control, mediated by NFkB. To this end, in a highly chemoresistant neuroblastoma PDX model, PRMT5 inhibition alone reduced PDX tumor growth and prolonged survival, associated with control of EMT-TFs, AKT and EGFR activity. Thus, our study has revealed a complex molecular interplay between PRMT5, AKT, EGFR, and NFkB, and their collective impact on EMT, both *in vitro* and *in vivo*.

We previously reported that PRMT5 mediated methylation of AKT is required for its subsequent phosphorylation^8^; further we now showed that PRMT5 directly regulates EGFR transcription. This dual regulatory role positions PRMT5 as a central mediator in neuroblastoma, capable of modulating multiple critical pathways driving tumorigenesis simultaneously. The independent regulation of AKT and EGFR by PRMT5 emphasizes the complexity of its function in cancer biology and suggests that its inhibition could have multifaceted therapeutic effects. In a neuroblastoma PDX model, PRMT5 inhibition reduced PDX tumor growth and prolonged survival. These results are consistent with the analysis of PRMT5 levels in neuroblastoma patients, where 95% of patients with the lowest quartile of PRMT5 were alive after 16 years but about 70% of patients with highest quartile of PRMT5 levels died within 4 years after diagnosis. Further, we identified NFkB as a downstream effector of AKT and EGFR in regulating EMT transcription factors in neuroblastoma. Given the well-documented role of NFkB in inflammation, cell survival, and proliferation, and recent reports of mesenchymal-like neuroblastoma cells displaying an inflammatory state^43, 44^, it is likely that PRMT5-dependent signaling pathways may also influence the tumor microenvironment, potentially impacting immune evasion and therapy resistance. However, since our studies were conducted in immunocompromised models, these results indicate that PRMT5 may be a viable therapeutic target and suggest that disrupting a PRMT5-dependent AKT-EGFR-NFkB signaling networks may be an effective strategy in treating high-risk neuroblastoma.

One of the most critical insights from our study is the elucidation of complementary loop between AKT and EGFR, each of which plays a significant role in reinforcing the EMT process downstream of PRMT5 activity. We showed that the reduced expression of EMT transcription factors upon PRMT5 inhibitor treatment could be rescued either by AKT or EGFR alone. This suggests that while PRMT5 inhibition impacts both AKT and EGFR pathways, these pathways compensate for each other, thereby sustaining the EMT process. This finding has profound implications for therapeutic strategies targeting neuroblastoma. The ability of AKT or EGFR to individually rescue EMT in the context of PRMT5 inhibition highlights the insufficiency of targeting either pathway alone. It underscores the necessity of a more general approach in therapy design, one that considers the interdependency and compensatory mechanisms of these signaling pathways. Therefore, inhibitors targeting solely AKT or EGFR may not be effective due to this compensatory regulation. Therefore, our results advocate for combined therapeutic approaches that simultaneously target both the AKT and EGFR pathways, such as PRMT5 inhibition, to effectively disrupt the EMT process and impede tumor progression.

## Methods

### Chemicals

PRMT5 methyltransferase activity inhibitor GSK3203591 was purchased from Cayman Chemical (cat#1616391-87-7). AKT inhibitor MK-2206 (cat# 11593), EGFR inhibitor Erlotinib (cat# 10483), and NFκB inhibitor Bay-117082 (cat# 10010266) were purchased from Cayman Chemical. All chemicals were dissolved in 100% DMSO as stock solution and subsequently diluted in phosphate-buffered saline or growth media to working solution.

### Cell culture

CHLA-20, SK-N-BE (2), and NGP cell lines were cultured as previously described^8^. For combination treatment, cells were cultured in the presence of 100 nM GSK3203591 for 6 days before exposure to 10 nM MK-2206 or 1 µM Erlotinib overnight.

### Plasmids

Human pCDNA3/Flag-HA-AKT1, Plncx/myr-HA-AKT1, and pCDNA6A/EGFR were purchased from Addgene as described previously^8^. The FLAG-PRMT5 and FLAG-PRMT5 (G367A,R368A) double mutant were from Dr. Said Sif (Qatar University) as previous described^45^. Transfections were performed as previously described^8^.

### RNA-seq

CHLA-20 and NGP cells were treated with vehicle or IC50 dose of GSK591 for six days. Dead cells were removed by Dead Cell Removal Kit (Miltenyi). Total RNA was extracted from live cells using Trizol reagent (Invitrogen) and purified by Direct-zol RNA Purification Kit (Zymo). RNA integrity was assessed using the RNA Nano 6000 Assay Kit of the Bioanalyzer 2100 system (Agilent Technologies, CA, USA). The mRNA enrichment, mRNA fragmentation, second-strand cDNA synthesis, size selection, PCR amplification, and sequencing were performed by BGI using DNBseq sequencing technology (One Broadway, Cambridge, MA 7 02142). RNA-seq reads from each sample were aligned using STAR against the GRCh38/hg38 human reference genome^46^. DESeq2 was used to identify differentially expressed genes using raw read counts quantified by htseq-count with GENCODE V29 gene annotation. GO term enrichment on differentially expressed genes was performed using DAVID^47–50^. Gene set enrichment analysis (GSEA) for differentially expressed genes was performed using pre-ranked gene lists ordered by −log10 (P value) multiplied by +1 for upregulation or −1 for downregulation.

### Cell viability assay

Cells were plated in 96-well plates and then maintained in the presence of a vehicle or various combinations of treatment for 72 hours. Cell viability was determined by MTS assay as previously reported^8^. The absorbance at OD490 was measured with a Synergy H4 Hybrid microplate reader (Bio Tek, Winooski, VT) using Gen5 software.

### Western blot

Tissue or cells were washed with cold PBS and lysed in RIPA buffer containing a protease and phosphatase inhibitor cocktail (Roche Diagnostics, Indianapolis, IN, USA). Proteins were analyzed by SDS-PAGE, followed by immunoblotting with indicated primary antibodies. Blots were incubated with horseradish peroxidase-conjugated anti-mouse or anti-rabbit IgG antibodies, and proteins were detected by enhanced chemiluminescence (ThermoFisher Scientific Inc., cat#7074). Antibodies used in this study are listed in **Supplementary Table 1**.

### Immunoprecipitation (IP)

Total proteins were extracted from cultured NB cells using RIPA buffer supplemented with protease and phosphatase inhibitor cocktail (Roche). The cell lysates were pre-cleared with protein A or protein G beads (Santa Cruz Biotechnology, cat#sc-20001, cat#sc-2002) at 4°C for 1 h prior to incubation with 5 μg primary antibody or isotype IgG at 4°C for overnight. Magnetic protein A/G beads (Millipore, cat# LSKMAGAG10) were added to the cell extracts and incubated at 4°C for 1 h. Then the beads were washed three times with washing buffer (100 mM NaCl, 50 mM Tris pH 7.5, 0.1% NP-40, 3% glycerol, 100 mM phenylmethylsulfonyl fluoride). The proteins bound to beads were eluted by adding 2x Laemmli sample buffer and heated at 95°C for 5 min. Eluted proteins were analyzed by western blotting.

### Cell invasion assay

*In vitro* invasion assay was performed as previously reported (NC). Briefly, cell culture inserts with 8 µm pore size (BD Biosciences, MA, USA) were coated with 50 μL extracellular matrix (ECM) from Engelbreth-Holm-Swarm murine sarcoma (Millipore Sigma, cat#E1270) and then hydrolyzed at 37°C for 30 min. Cells were detached and suspended at 5×10^4^ cells/ml in serum-free media, and two hundred microliter cell suspension was added onto the upper surface of the insert. Seven hundred and fifty microliters of growth media were added to the bottom chamber. After 15 h incubation, cells grown on the top surface of the insert were removed with cotton swabs. Cells that invaded the opposite surface of the insert were washed with PBS, fixed in 4% formaldehyde for 5 min, followed by methanol 100% for 20 min, and then stained with 0.25 % crystal violet for 10 min. Photographs were taken under a light microscope and quantified using Image J software. The percentage of cells of invasion was normalized to the equal number of cells plated in 24-well plate cultured for the same time period.

### Patient-derived xenograft mouse model

NOD/SCID mice were used in this study. Ten million COG-N-623X PDX cells (Children’s Oncology Group) were suspended in 100 µL PBS and mixed with 100 µL Matrigel were implanted in the upper back of an NSG mouse subcutaneously. Tumor size was measured by digital caliper three times a week, and tumor volumes were estimated using the modified ellipsoid formula V = ab2/2 (a>b; a, length; b, width) as previously. When tumor volume reached 300-500 mm3, mice were randomized into either a “vehicle control” group (0.5 methylcellulose, Sigma-Aldrich #M0430) or a PRMT5 inhibitor “GSK595 treatment” group (50 mg/kg). Animals were treated for three weeks, with vehicle or 500 mg/kg GSK595 by oral gavage twice daily. The body weight of mice was monitored weekly. Mice were euthanized once tumor volume reached 2,000 mm^3^ or if moribund from toxicity or illness. All procedures were approved by the Institutional Animal Care and Utilization Committee (IACUC) at UMass Medical School.

### Statistical Analyses

All quantitative data points represent the mean of three independent experiments performed in at least triplicates with Standard error of the mean (S.E.M). Unless indicated in the figure legend, statistical analysis was performed using a two-tailed, unpaired t-test, or survival analysis (GraphPad Software, Inc., La Jolla, CA).

## Acknowledgements.

This work was supported by Dean’s Research Fund (University of Massachusetts Medical School) and Hyundai Scholar Hope Grant (2019, 2022) (Hyundai Hope on Wheels Foundation). A.D.D. is supported by NIH grant K08-CA245251, the Rally Foundation for Childhood Cancer Research, Hyundai Hope on Wheels Foundation and V Foundation for Cancer Research. A.D.D, N.A.M.S and M.A.M.N are supported by the American Lebanese Syrian Associated Charities (ALSAC).

## Author contributions

L.H., R. V., V. O., S.G., X.X., N.A.M.S and M.A.M.N designed and performed the experiments, and analyzed the data. Z.X. performed bioinformatic analyses. W.G. performed *in vivo* PDX experiments. A.D, T.F, and W. Q designed the experiments and analyzed the data. W.Q. conceived the project and prepared the manuscript.

