## Supplemental information for "PRMT5 orchestrates EGFR and AKT networks to activate NFκB and promote EMT"

**Supplemental Figure1. Inhibiting FGFR4 and NGFR did not affect the expression of EMT-TFs.**

CHLA20 and NGP cells were treated with DMSO or 100 nM GSK591, 1μM of Erlotinib, 10 μM of NGFR inhibitor PD 90780, 10 μM of FGFR4 inhibitor FGFR4-IN-1 for 24 hours. Cells were collected and whole cell lysate was extracted for western blot analysis.

**Supplemental Figure2. PRMT5 does not methylate NFκB in neuroblastoma.**

The p65 subunit of NFκB was immunoprecipitated from CHLA20 and NGP cells were treated with DMSO or 100 nM GSK591, then the SDMA of p65 as well as pull downed p65 were examined by Western blot (upper panel). The expression of p65, SDMA, and Actin was analyzed in 10% of input from each sample by Western blot.

### Extended Data Figure 1

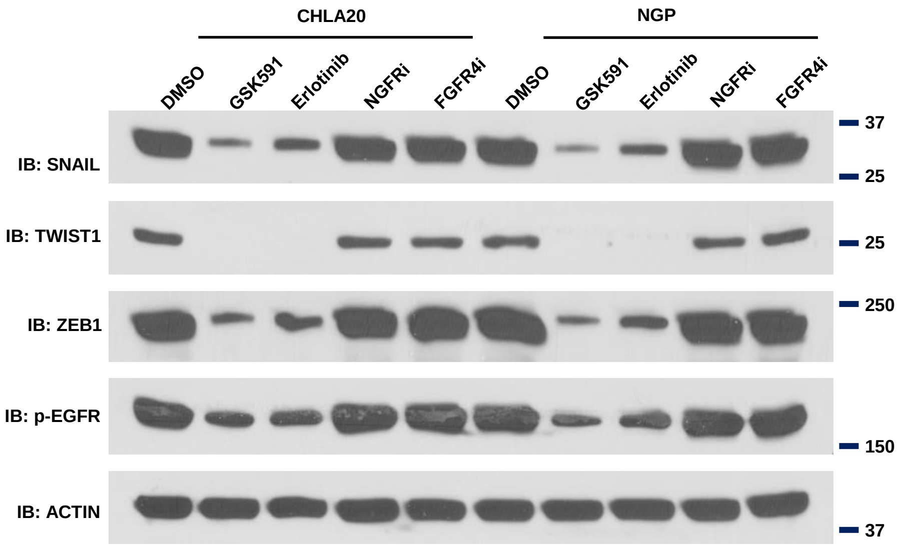

### Extended Data Figure 2

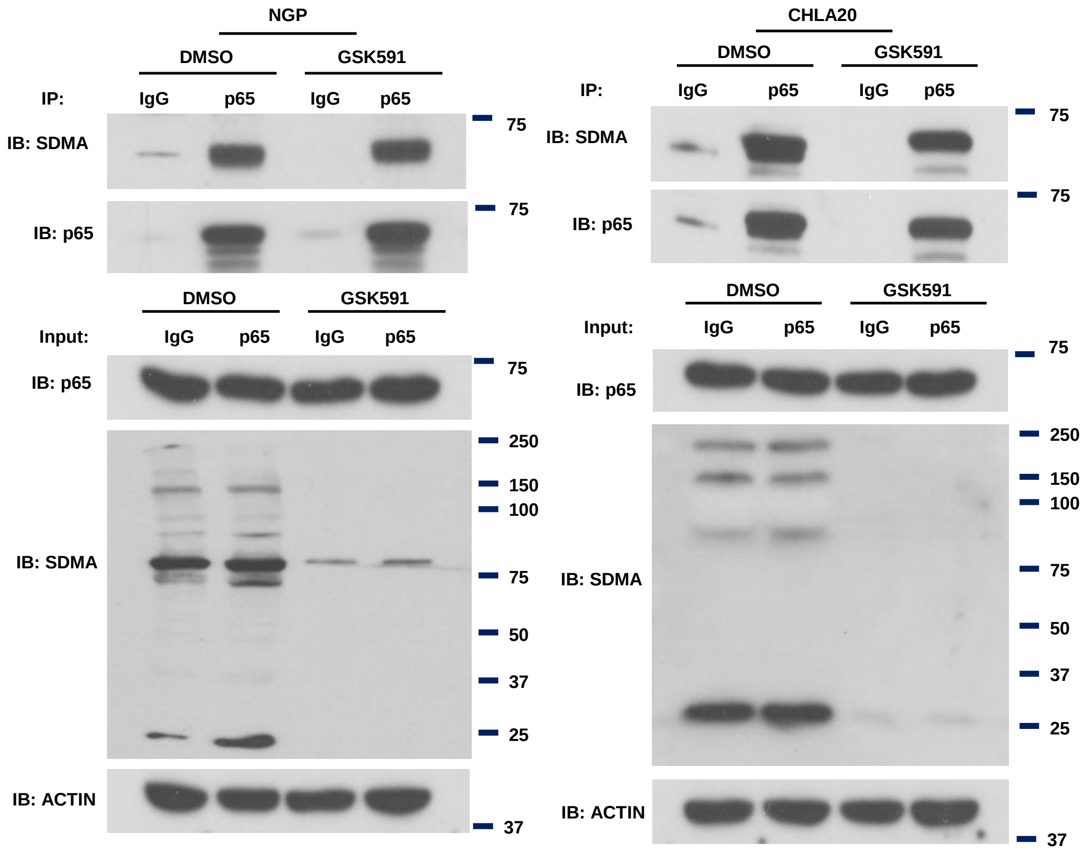

**Table 1. Antibodies used in this study**

| Name | Vendor | Catalog Number |
|---|---|---|
| AKT | Cell Signaling | 9272 |
| phospho-AKT Ser 473 | Cell Signaling | 4060 |
| Phospho-(Ser/Thr) Akt Substrate | Cell Signaling | 9611 |
| EGFR | Cell Signaling | 4267 |
| phospho-EGFR Tyr 1068 | Cell Signaling | 3777 |
| FGFR4 | Cell Signaling | 8562 |
| Flag | Sigma-Aldrich | F7425 |
| anti-HA-Peroxidase | Sigma-Aldrich | 12013819001 |
| phospho-Tyr-1000 | Cell Signaling | 8954 |
| p75NTR | Cell Signaling | 8238 |
| PRMT5 | Santa Cruz | sc-376937 |
| ZEB1 | Cell Signaling | 70512 |
| SNAIL | Cell Signaling | 3879 |
| TWIST1 | Cell Signaling | 46702 |
| RNA Pol II CTD | ThermoFisher | 49-1033 |
| PRMT5 (ChIP grade) | ThermoFisher | PA5-78323 |
| NF-κB p65 (D14E12) | Cell Signaling | 8242 |
| Phospho-NF-κB p65 (Ser536) (93H1) | Cell Signaling | 3033 |
| Histone H3R8 Dimethyl Symmetric | Epigentek | A-3706 |
| Histone H4R3 Dimethyl Symmetric | Epigentek | A-3718 |
| beta Actin | Santa Cruz | sc-47778 |
| anti-GAPDH-Peroxidase | Sigma-Aldrich | G9295 |
| Mouse anti-rabbit IgG-HRP | Santa Cruz | sc-2357 |
| Mouse IgGκ BP-HRP | Santa Cruz | sc-516102 |
| Alexa Fluor 488-conjugated goat anti-rabbit | ThermoFisher | A32731 |
| Alexa Fluor 596-conjugated goat anti-mouse | ThermoFisher | |
